# Cooperative dynamics of PARP-1 Zinc-finger domains in the detection of DNA single-strand breaks

**DOI:** 10.1101/2024.06.09.598140

**Authors:** Parvathy A. P. Sarma, Corinne Abbadie, Fabrizio Cleri

**Author notes:** (corr. author).

## Abstract

The DNA single-strand break (SSB) repair pathway is initiated by the multifunctional enzyme PARP-1, which recognizes the broken DNA ends by its two zinc-finger domains, Zn1 and Zn2. Despite a number of experiments performed with different DNA configurations and reduced fragments of PARP-1, many details of this interaction that is crucial to the correct initiation of the repair chain are still unclear. We performed Molecular Dynamics (MD) computer simulations of the interaction between the Zn1/Zn2 domains of PARP-1 and a DNA hairpin including a missing nucleotide to simulate the presence of an SSB, a construct used in recent experiments. The role of Zn1 and Zn2 interacting with the SSB ends is studied in detail, both independently and cooperatively. We also explored, PARP-1 operating as a dimer, with the two Zn-fingers coming from two separate copies of the enzyme. By an extensive set of all-atom molecular simulations employing state-of-the art force fields, assisted by empirical docking and free-energy calculations, we conclude that the particular conformation of the DNA hairpin can indeed spontaneously open up by thermal fluctuations, up to extremely kinked deformations. However, such extreme localized deformations are rarely observed in free, long DNA fragments. Protein side-loops make contact with the DNA hairpin grooves, and help Zn2 to penetrate deep in the SSB gap. In this way, Zn2 can interact with the nucleotides opposite to the missing base. OVerall, Zn1 plays a secondary role: the crucial factor for the interaction is the relative arrangement of the Zn1/Zn2 couple, and their mutual orientation with respect to the 3^′^ and 5^′^ SSB end terminals. This helps to obtain an early interacting configuration, which ultimately leads to molecular PARP-1-DNA structures similar to those observed experimentally. Such findings represent an important step toward defining the detailed function of PARP-1 in the early stages of SSB recognition.

## 1 Introduction

Ionizing radiations induce damage in the molecular components of the cell, chiefly including DNA, the most critical component for cell survival and proliferation^1–3^. The most lethal damage is usually considered the double-strand break (DSB), defined as two breaks in the sugar-phosphate backbone on each DNA strand separated by less than ten base-pairs. The signalization and repair of such molecular damage involve the activation of the DNA damage response (DDR) pathway with downstream activation of the tumor suppressor TP53. This can lead to a transitory cell cycle arrest, thus favoring DNA repair, or induce apoptosis. Often considered less dangerous, DNA single-strand breaks (SSBs) are one of the most frequent forms of DNA damage. SSBs can be produced directly from oxidative damage, as well as intermediates in other DNA repair pathways. At the molecular scale, they are seen as a break in the sugar-phosphate backbone involving only one DNA strand, often accompanied by a nucleotide loss and displaying abnormal 5^′^ and 3^′^ ends^4^. Although being repaired very efficiently, even a small fraction of unrepaired SSBs can compromise DNA replication and transcription, leading to genome instability, frequently associated with cancers and neurodegenerative disorders^5^. For example, we demonstrated in a recent work^6^ that therapeutic irradiation by high-energy photon beams can induce an abnormal accumulation of SSBs in fibroblasts exposed to out-of-field dose. The affected cells consequently enter senescence and may subsequently lead to secondary cancers upon escaping from their dormant state with residual damage.

The SSB repair (SSBR) pathway involves a first step of break detection initiated mainly by the poly(ADP)ribose polymerase-1 (PARP-1) and to a lesser extent by PARP-2. Once activated by interacting with the DNA strand break, PARP-1 starts to synthesize long branched chains of poly(ADP)ribose (PAR) using NAD^+^ as substrate. The accumulated PAR chains, either free or linked to proteins like histones (PARylation), or on PARP-1 itself (autoPARylation), favor the recruitment of the X-Ray Repair Cross-Complementing Group-1 (XRCC1) scaffold protein. XRCC1 is then phosphorylated and recruits the following repair enzymes such as the polynucleotide kinase phosphatase (PNKP) and polymerase-*β*. Notably, these last steps are common with the base-exchange DNA repair pathway (BER)^7^.

PARP-1 (Figure 1a) is a well studied member of PARP family. It is a chromatin-associated protein consisting of at least six functional domains: three DNA-binding Zinc-finger N-terminal domains named Zn1, Zn2 and Zn3; one BRCT domain; one WGR domain; and one catalytic C-terminal domain, including a helical subdomain (HD) (Figure 1c). When PARP-1 is not bound to DNA, Zn1 and Zn2 behave as flexible independent domains^8^. PARP-1 binding to damaged DNA activates a complex sequence of allosteric and cooperative effects between the different domains, which are not yet completely elucidated. Several studies were performed but using only fragments of the whole protein, often leading to divergent models, especially whether PARP-1 recognizes and binds to damaged DNA as a monomer or a dimer. Zn1 and Zn2 are known to specifically recognize DNA breaks (notably, both SSB and DSB). Zn1 from one PARP-1 copy may also cooperate with Zn2 from another PARP-1 protein to form a dimeric module that specifically recognizes DNA breaks^9^. On the other hand, Zn3 mediates as an inter-domain contact and is required to confer with PARP-1 to regulate chromatin structure^10^. The BRCT domain acts also as a DNA binding domain, but of low affinity, and is able to bind only intact DNA without concomitant catalytic activation. The BRCT-DNA interaction mediates DNA intra-strand transfer of PARP-1 (the so-called “monkey-bar mechanism”) that allows rapid movements of PARP-1 through the chromatin^11^. By analogy with PARP-2, it is assumed that the WGR domain of the sister protein domain of PARP-1 can bridge two nucleosomes with the broken DNA ends aligned in a position suitable for ligation. Such bridging induces structural changes in PARP-1 that signal the recognition of a DNA break to the catalytic domain of PARP-1. This promotes the recruitment of Histone PARylation factor 1 (HPF1) and subsequent activation of PARP-1, followed by licensing serine ADP-ribosylation of target proteins^12,13^. HD prevents effective NAD^+^-binding in the absence of an activation signal. However, after binding to damaged DNA, the autoinhibition is relieved, HD unfolds, and PARP-1 becomes able to bind NAD^+^, thus starting PARylation^14,15^.

**Figure 1.**
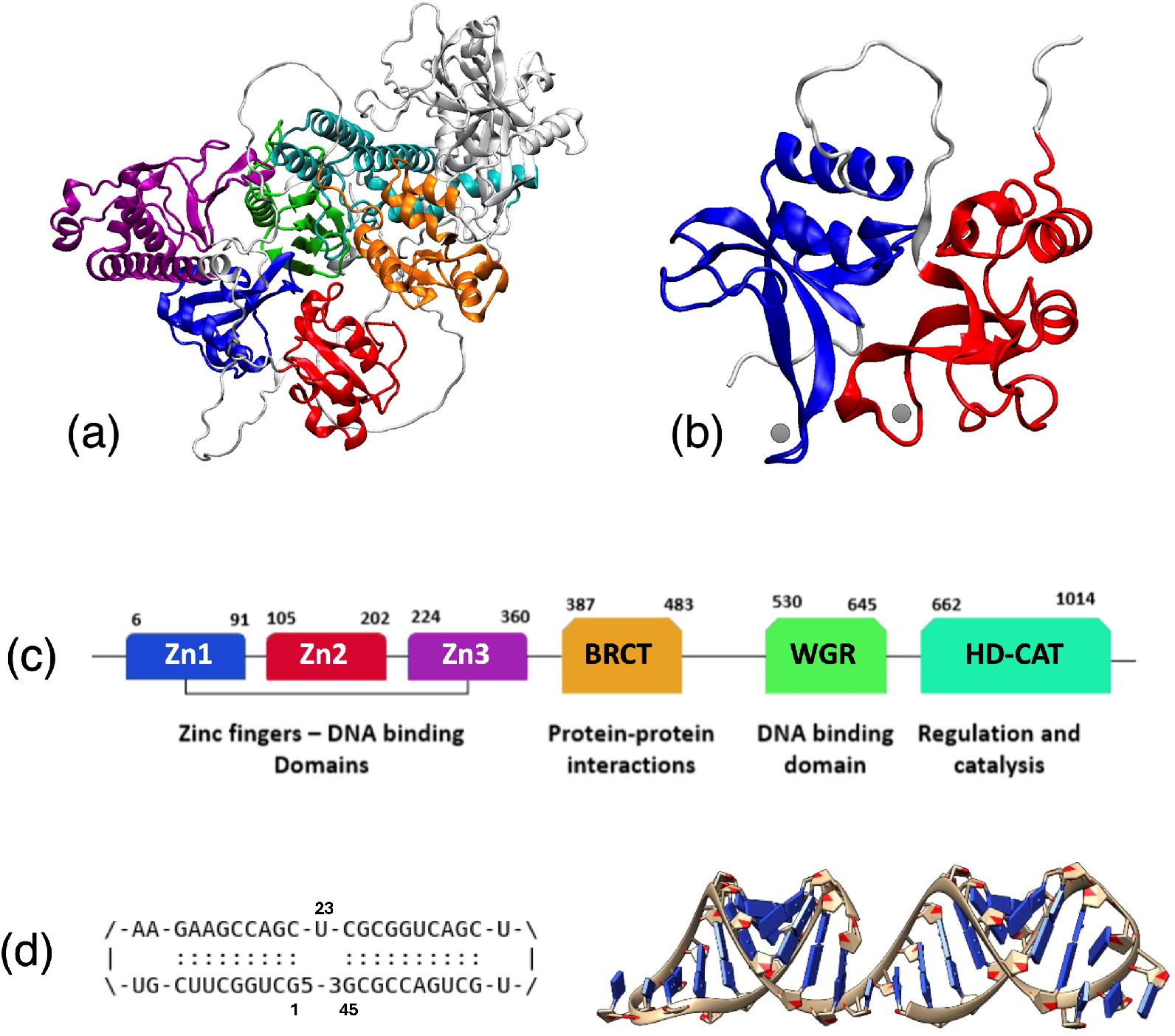
Molecular structures of the PARP-1 protein and of the DNA hairpin. (a) Whole structure as modelled by AlphaFold2. Domains: Zn1, blue; Zn2, red; Zn3, purple; BRCT, orange; WGR, green; PARP-*α*-helical, cyan. (b) Details of the two zinc-finger domains, Zn1 (blue) and Zn2 (red), that recognize and interact with the DNA single strand breaks. The position of the Zn atoms (grey spheres) is tentatively indicated. (c) Domain structure of PARP-1; the numbers above the diagram indicate the amino acid positions. (d) Blueprint of the RNA sequence used as input in ROAD along with the nucleotide positions 1,23,45, and the resulting 3D-structure of the straight DNA hairpin, with a missing nucleotide to simulate a single-strand break.

Some studies alread shed light on the structural features of Zn1 and Zn2 domains interacting with DNA breaks. In the earlier works, single PARP-1 fragments (with either Zn1 or Zn2) were co-crystallized with short dsDNA fragments, ideally mimicking DSBs with blunt ends^16^. The results showed that both Zn1 and Zn2 can make contact with the DNA ends through interaction of Phe44 and Lys151 respectively with the 5^′^ and 3^′^ ends of DNA preferentially. However, Zn2 showed a higher affinity than Zn1 in binding to DNA ends. A subsequent study, in which dsDNA terminated with overhangs at both ends were used^9^, and co-crystallized with separate Zn1 and Zn2 domains (ideally representing the N-terminals of two adjacent PARP-1 molecules), found that Zn2 interacts with *both* the 5^′^ and 3^′^ ends, tending to exclude Zn1 from the direct interaction but requiring the cooperation of both Zn fingers for the efficient tagging of the damaged DNA *in vivo*. Eustermann et al.^14^ reported a high-resolution structure of PARP-1 Zn domains in complex with an SSB. The authors used a hybrid nuclear magnetic resonance and X-Ray diffraction (NMR-XRD) technique to produce a detailed atomic structure of paired Zn1-Zn2 domains in interaction with a 45-nucleotide DNA hairpin containing a SSB. They showed how Zn2 initiates the recognition of the SSB by first binding to the 3^′^ end, followed by Zn1 (from the same PARP-1 monomer) making contact toward the 5^′^ end. This coupled action appears to lead to a drastic bending of the DNA about the SSB site, by ∼130°, an effect that was also observed in older experiments by electron microscopy^17^. Very recently, the same group also performed single-molecule Förster resonance energy transfer (smFRET) analysis on the same DNA-hairpin with SSB in interaction with Zn1 and Zn2^18^, showing that just the co-presence of Zn1-2 (even excluding the rest of the protein) is sufficient to bring about a kink in the DNA at the break site; it appears that Zn2 recognizes the damage by producing a small kink, and additional binding of Zn1 induces a sharper kink in the DNA.

Even though these structural studies helped in understanding many details of the binding of PARP-1 to damaged DNA and its consequent conformational changes, there is no clear-cut answer on the mechanism by which the change in conformation of DNA occurs.

Moreover, the initial steps of the damage recognition are not established, since the biochemical techniques used until now only give access to final-state conformations. In this work, we combined molecular docking and molecular dynamics (MD) simulations, to understand the pristine interactions of PARP-1 with the damaged DNA, and the dynamics of kinking-bending that leads to the stable binding between the enzyme and the damaged DNA site. We started from a DNA system similar to that used by Eustermann et.al^14^ and retrieved the kinked dumbbell shape already reported. Then, we also used a straight DNA hairpin loop, carrying the same sequence and defect, to investigate the early stages of recognition by Zn1 and Zn2, both separately and in tandem. A careful comparison was also performed between monomeric and putative dimeric interactions of PARP-1.

We conclude that the highly peculiar conformation of the 45-nucleotide long DNA hairpin with end-loops, used to model a SSB in the experiments, can spontaneously open up by thermal fluctuations, up to extremely kinked deformations, with the two DNA arms flanking the SSB making angles as small as 90°-100°. On the other hand, we rarely observed such extreme localized deformations in a free, long DNA sample with standard blunt ends.

An important feature of this peculiar construct is represented by the Zn1 and Zn2 side loops, respectively centered at Arg-18 and Arg-122, which can make contact with the DNA hairpin grooves and help Zn2 to penetrate deep in the SSB gap, up to reaching the opposite strand where it interacts with the unpaired nucleotide facing the SSB. Zn1 seems to play a secondary role; however, the relative arrangement of the Zn1/Zn2 couple, and their mutual orientation with respect to the 3^′^/5^′^ ends is key to obtain a properly interacting configuration. This, ultimately leads to the formation of molecular PARP-1-DNA structures compatible with those deduced from biochemical and biophysical techniques. However, simulations also suggest that the *invivo* recognition mechanisms could follow somewhat different kinetic paths, compared to the very special hairpin construct.

## 2 Methods

To elucidate the molecular interactions between the zinc-finger domains of PARP-1 and the DNA damaged by a SSB, we designed a complete simulation protocol, starting with (i) the design of the atomic structure of the DNA and protein fragments, then (ii) docking of the rigid fragments (protein to the DNA) with an approximate free-energy functional, followed by (iii) molecular dynamics simulations and analysis with a state-of-the-art empirical force field, and finally (iv) comparison with the available experimental results.

In the following we use the same DNA hairpin including an SSB as a reference receptor, that was the source structure used in experiments^14,18^. Since, the reported NMR experimental structure only presents a final state with a strongly deformed hairpin, as a result of the interaction with the protein, we defined also a straight DNA hairpin as another initial condition for the simulations, with the same nucleotide sequence and 2D contacts. By using this at the initial configuration, we could also follow the process of SSB recognition from the earliest stage of the interaction.

To design the DNA hairpin, we used the RNA-Origami Automated Design tool (ROAD), a computer-aided design software, to build automated 3D-structures of RNA^19^. We submitted the blueprint of a 45 nucleotide long RNA, with the same experimental DNA sequence of Ref.^14^, and simply substituted Uracil (U) with Thymine (T) using CHIMERA^20^. The resulting DNA structure comprises of missing nucleotide at the center, mimicking an SSB (Figure 1d), flanked at 3^′^ and 5^′^ by Guanine in positions 1 and 45, and an unpaired Thymine in position 23 facing the missing SSB base. We also made sure that the DNA forms a closed loops with 2 unpaired nucleotides in positions 11, 34 and 35, at both hairpin ends. This DNA structure was then utilized as a second starting configuration, next to the experimental one, to elucidate the first steps of the PARP-1 Zn-finger interactions with the DNA. While there could be minor differences between RNA 3D structure and the putative dsDNA, based on our previous studies following this same protocol^21^ we are confident that the subsequent equilibration by MD at finite temperature can wash out any structural instability, and produces a stable DNA hairpin. We used CHIMERA^20^ to design a long stretch of free DNA with 200 nucleotides, with a missing base in one strand at position 101 to represent the SSB. Like for the hairpin, the break in this long DNA is also flanked by Guanine on both 5^′^ − 3^′^ sides, along with Thymine facing the missing nucleotide.

The structure of the Zn1-Zn2 finger region of PARP-1 was taken from the experimental NMR data^14^, deposited in the RCSB PDB archive under the entry **2N8A**. For the sake of comparison, we also took the corresponding fragment from the AlphaFold2 repository, entry **P09874** (see Figure 1)^22,23^. The structure obtained from **2N8A** consists mainly of the Zn1-Zn2 fragment of PARP-1 (residues 1-201). AlphaFold2 instead predicted the entire structure of PARP-1 with all its domains: in the AlphaFold2 model, all major domains of PARP-1 are comparable to the corresponding solution structure obtained from NMR studies within a reasonable confidence interval, except the disordered linker stretch of 13 amino acids connecting the Zn1 and Zn2 domains. Docking of proteins and nucleic acids is still a rapidly evolving field of research. We tested different methods available as online server platforms: HADDOCK^24,25^, HDOCK^26^ and the very recent pyDockDNA^27^. In our analyses, only HADDOCK was capable of providing consistent results comparable to the available experimental information. This is evident in both the straight DNA hairpin and the bent DNA extracted from the experimental NMR structure, especially when assessed against the Zn1-Zn2 fragment of PARP-1. HADDOCK bootstrap parameters were chosen such that we could typically aim at the regions including the trademark interactions between Zn1/Zn2 and the SSB^14^. Best-fit clusters of candidate high-scoring poses were taken from the docking process.

Furthermore, we performed interpolation morphing between the straight and bent DNA conformations, using CHIMERA. Such morphing simulations provided a preliminary look at how the protein and DNA might change their conformations during the SSB recognition process. From the first step, it seems that no steric clashes are prohibiting the rearrangement towards the final (experimental) conformation. However, the results of docking displayed a large variety of candidate conformations, also including inverted contacts, that is Zn1 close to the 3^′^ end and Zn2 close to the 5^′^ end of the hairpin, contrary to the experimental evidence. Hence, starting from the docked configuration, we also tried to improve the contacts by refinement, adjusting the contacts by minimal rigid-body translations and rotations, with subsequently relaxing and cyclic annealing by MD of the “post-docked” configurations. These adjusted conformations will be also used in the following as candidate starting structures for the SSB recognition.

All the MD simulations and most data analysis, were performed with the GROMACS 2020.4 package^28,29^. The Amber14 force-field was used in the simulations^30^, with extra parameters for Zn ion interaction from Macchiagodena et al.^31^; notably, Amber14 already includes the lastest PARM-BCS1 extension for nucleic acids^32^. To ensure the primary stability of the post-docked configurations, we first performed energy-minimization using a steepest-descent algorithm, followed by {NVT}-{NPT} equilibration for 150ps, to bring the systems at 310K and 1 bar pressure. The ensembles of DNA-proteins were solvated in water boxes of typical size 15×15×15 nm^3^, with about 106,500 water molecules, along with Na^+^ and Cl^−^ ions for neutralizing the system charge and maintaining a physiological 0.1M salt concentration. In some instances we also performed rapid annealing and quenching cycles, to improve the mutual positions of protein and DNA, for example after a manual adjustment necessary to remove a steric clash. Each cycle was typically performed by ramping up the temperature from 310K to 400K, and then back to 310K, in steps of 10K for 500 ps each. Usually, during the annealing cycle the DNA was frozen in its conformation and only the protein was left free to adjust.

Coulomb forces were summed with particle-mesh Ewald sum (PME), using a real-space cutoff of 1.2 nm, equal to the cut-off radius of shifted Van der Waals potentials. We used rigid bonds for the water molecules, which allowed integration of the equations of motion with a time step of 2 fs for the thermal equilibration phases, and 1 or 2 fs for production runs.

Trajectory clustering analysis was performed using a special subprogram of GROMACS. The MD simulation stores ‘frames’ containing the positions, velocities and forces of all particles in the system, at prescribed intervals (typically every 10 to 50 ps, or longer: a 1 *µ*s-long MD run can store as much as 100,000 frames of about 2Mbytes each, resulting in data files with size of hundreds of Gbytes); the subprogram calculates a matrix of root-mean-squared displacements (RMSD) between each pair of frames, by comparing the positions of each atom in the pair; then, RMSD values are grouped according to a cut-off criterion, and clusters of similar frames, typically separated by a small enough RMSD, are detected. The cluster(s) with the highest number of members, and those which are more frequently sampled over the entire simulation time, provide an indication of the ‘best-average’ molecular configurations, thus allowing to bypass the noise of the short-time atomic fluctuations.

Free energy analysis and contact surface calculations were estimated by the PDBePisa web-based utility^33^, by extracting a few selected configurations in PDB format from the MD trajectory. The resulting values are approximated, however they can provide a qualitative appraisal of the binding energies involved in the protein-DNA interaction.

## 3 Results

Strand breaks in DNA are the result of a complex sequence of biochemical modifications, starting from the simple breaking of a bond and proceeding in several steps with the progressive disruption and final removal of the damaged nucleotide(s). Hence it is crucial to understand how, and at which step, the early SSB detection is activated by the sensor proteins that continuously search the DNA for structural anomalies. In the following, we used the DNA hairpin of Ref.[14] as the reference system to study the interaction of PARP-1, both in monomeric and dimeric form, with a model-SSB represented as a whole missing nucleotide in the center of the DNA hairpin. Such a structure is representative of the direct damage to the DNA, as can be obtained as a result of the rapid chemical evolution of the initial bond-breaking typically produced at the C4^′^ or C5^′^ carbon by radical attack.

### 3.1 Stability of the isolated DNA hairpin including a SSB

We first tested the thermal and mechanical stability of the model DNA hairpin including a simulated single-strand break. In particular, we wanted to verify whether the DNA fragment could bend spontaneously into the experimentally observed shapes^14^, under the simulated thermodsynamic conditions, and without direct participation from PARP-1. For this purpose, we performed 2*µ*s-long MD simulations for three variations of the model DNA: **(1)** the 45-nt hairpin from Ref.[14] with a missing nucleotide at its center representing the SSB, with the same nucleotide sequence and terminations as the experimental structure (5^′^-phosphate and 3^′^-OH, Figure 1d); **(2)** the same hairpin as in **1** but with hydroxyl terminations on both 5^′^ and 3^′^ ends, a configuration that may be representative of the SSB gap after the pol-*β* removal of 5^′^-dRP, and could be interesting to compare the effect of a wider opening of the gap; **(3)** a long stretch of free DNA with a missing nucleotide at one strand, locally reproducing the same SSB gap in the hairpin.

The MD simulation of the isolated hairpin **1**, shows that this DNA structure does spontaneously open up at the level of the SSB, and can bend substantially, over very short time frames. Even though the DNA largely fluctuates in the course of the simulation (Figure S1-a,b), the average stable structure after cluster analysis (cutoff 0.6 nm, average RMSD 0.49 nm) remains comparable to the initial straight configuration (Fig.S1-c). We measured the gap opening at the SSB by taking the distance *d*_*g*_ between the C4^′^ atoms of the guanine 1 and 45, respectively at the 5^′^ and 3^′^ terminals. The bending angle *θ*_*b*_ is measured by the relative position of the P atoms of the nucleotides 12, 24 and 34, at the two ends and approximate center of the DNA. Figure 2a,d displays the plot of these values for the 2-*µ*s trajectories, the blue, red and black colors referring to the structures **1**,**2**,**3**, respectively.

**Figure 2.**
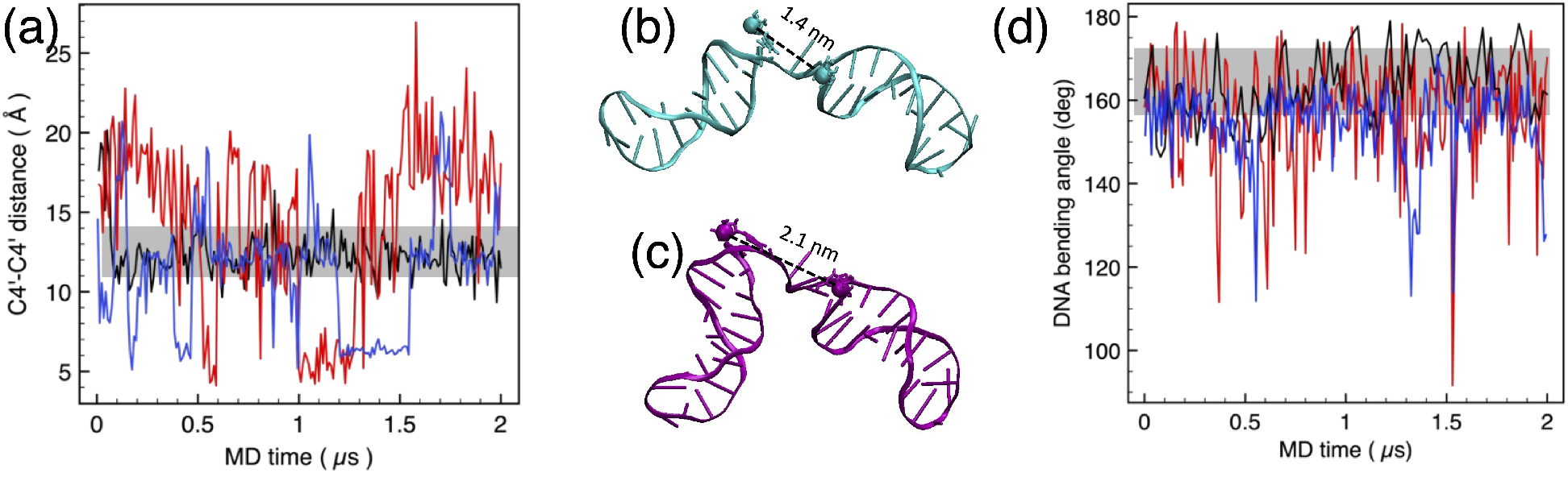
(a) SSB opening, measured as the distance between the C4’ atoms of the 5^′^ and 3^′^ guanines: straight long DNA (black), 45-nt hairpin with 5^′^-dRP (blue), and with 5^′^-OH (red). (b,c) Sample snapshots of the hairpin configuration at times 0.5 and 1.77 *µ*s, showing typical frames of the intermediate (b) and wide-open conformation (c). (d) Bending angle, measured between the P atoms of nucleotides 12-24-34 (center and loop ends). To facilitate the comparison, the grey shaded areas highlight 95% of the fluctuation for the straight long DNA (black).

The SSB gap displays large jumps between three different states: a “wide open” at *d*_*g*_ ≥2 nm, an intermediate one at *d*_*g*_ ∼1.5 nm, and a sort of “closed” state with a gap distance *d*_*g*_=0.5-0.7 nm. The DNA bending angle experiences large fluctuations between *θ*_*b*_=110° and 160° (Fig. 2d), with instantaneous peaks at even smaller angles. The coupling between opening and bending indicates large fluctuations accompanied by a (less visible) twisting of the compact DNA structure. In particular, the peaks appearing at times *t*=1.58, 1.77, 1.83 and 1.87 *µ*s, are correlated with the corresponding jumps in the RMSD (Fig.S1-b); the snapshots of the DNA hairpin in Figure 2b,c show an extensive bending and twisting of the structure in both the intermediate and wide-open configurations.

The comparison of structure **2** with dephosphorylated 5^′^ end (see red plots in Figure 2) is entirely similar to the structure **1**, and displays an even wider range of fluctuations both in the opening and in the bending angle. The RMSF-by-atom shows a distinct periodicity, with larger values of about 0.6 nm for the two 5^′^ and 3^′^ ends, the nucleotides 11-12 and 34-35 at the loops (up to 0.8 nm for the 11-nt), and the unpaired nucleotide 23 facing the gap. This indicates that the two end loops in this particular experimental construct could help the spontaneous opening-up of DNA, both before and during the interaction with the PARP-1 protein.

For the sake of comparison, we extended the same stability test to the free DNA structure **3** that could be representative, e.g., of a nucleosome linker, or a transcriptionally exposed chromatin stretch. We chose a DNA length of 200 bp (about 70 nm) which is longer than the typical DNA persistence length^34,35^. This structure includes a gap at the centre of the double strand (resembling an SSB) and no end loops. Evidently, such a long DNA displays long-wavelength fluctuations at the simulation temperature of 310K, as it can be observed by the time plot of the RMSD and the RMSF-by-atom, which fluctuate about values of several nm. However, when we looked at the gap opening and the DNA bending angle (Figure 2a,d) an important difference is observed between the free DNA (black plots) with respect to the same quantities measured on the 45-nt hairpin (blue and red plots). The gap opening in the free DNA remains at a constant value of about 1.2 nm with a small dispersion of *±*0.1 nm. The average bending angle in the free DNA fluctuates between *θ*_*b*_=160-175°.

Such results suggest that the large opening of the SSB gap apparently necessary to accommodate PARP-1 in the experiments using the DNA hairpin^14^, could be due to - or at least be substantially helped by - its spontaneous thermal fluctuations. However, our results indicate that the wild bending and gap opening of the hairpin seems confined to shorter (tens of ps) times, while the average values remain close to the initial conformation. By contrast, it may be suggested that the *in-vivo* SSB conformations (being entropically constrained by the more rigid chromatin structure) could be more stable and display less frequent spontaneous openings, as suggested by the straight-DNA results.

### 3.2 Binding of PARP-1 monomer to DNA hairpin

We then proceeded to study the binding of PARP-1 to DNA containing a SSB. As detailed in the Methods section, we used molecular docking with empirical free-energy functional (HADDOCK) to identify the candidate interacting conformers, followed by finite-temperature MD with AMBER14+BSC1 force fields, to unveil the binding dynamics of PARP-1 and conformation fluctuations. As a first test to establish the validity and quality of the simulation protocol, we initially tried to retrieve the experimental interaction from Ref.^14^. For the sake of clarity, in the following we will indicate protein residues by their 3-letter code and residue number, and DNA nucleotides by a single letter-number pair.

#### 3.2.1 Retrieving the experimental structure

We generated two separate PDB structures by isolating the protein and the DNA hairpin from the RCSB PDB entry 2N8A, which were then reassembled by docking with HADDOCK. The active residues to bootstrap the docking were: the residues Phe-44 and Leu-151 of the protein (belonging to Zn1 and Zn2 respectively), and the nucleotides G1 (5^′^ end of the hairpin), G45 (3^′^ end), plus the T23 (unpaired base facing the SSB gap) of the DNA. This resulted in docked clusters having some of the interactions that were already established in the experimental NMR structure. Then, we selected the best docking pose based on the RMSD value as initial configuration, and performed a 200-ns MD simulation at T=310K (Figure 3a). The resulting probability distribution of the 5^′^-3^′^ distance (i.e. the SSB gap), as measured by the distance between the C4^′^ atoms of G1-G45, (Figure 3b, black line) has a peak at *d*_*g*_ ≃3.1 nm, compared to 3.02 nm for the experimental NMR structure^18^. The distribution of the DNA bending angle, shown in Figure 3c (black line), extends between about *θ*_*b*_=80-160°, peaked at about 135°, and rather skewed toward the lower values. By comparison, the distribution of experimental values obtained by FRET^18^ is nearly Gaussian, peaked at 90 ± 14°.

**Figure 3.**
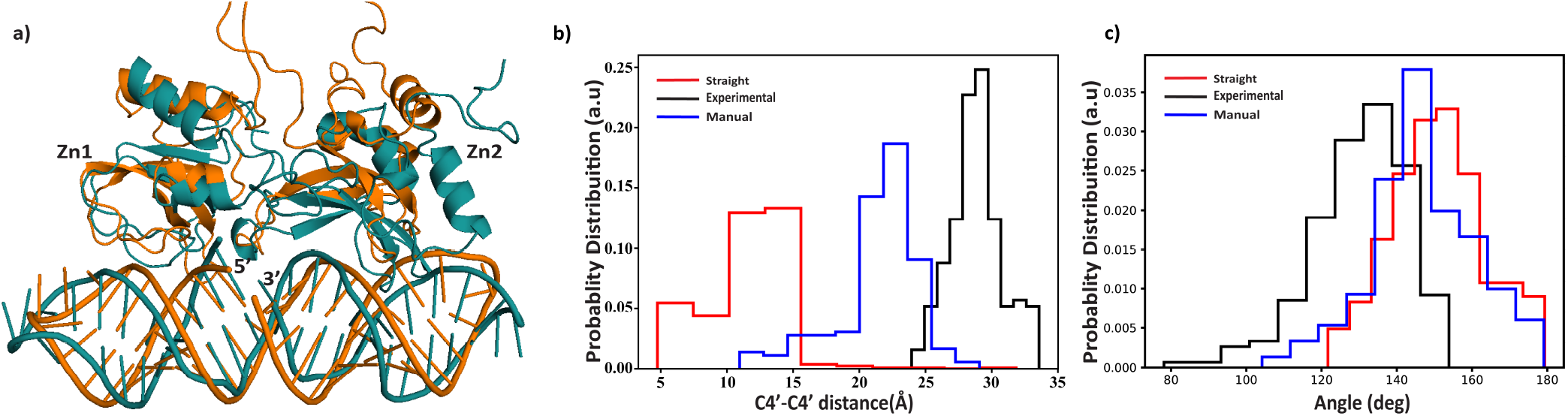
(a) Straight hairpin before (orange) and after (cyan) 500ns simulation. (b) Distance between the C4^′^ atoms of the 3^′^ − 5^′^ ends: DNA starting from experimental structure (black) and free PARP-1 docking. DNA starting from the straight hairpin (red) and free PARP-1 docking. DNA starting from the straight hairpin (blue) and manual fitting of PARP-1. (c) Distribution of DNA hairpin bending angle measured between the P atoms of nucleotides 12-24-34; same color code as in (b).

In the simulated complex, we find a very close proximity between Leu-151 from Zn2, and the DNA nucleotide G45 at the 3^′^ end; and the Phe-44 from Zn1 in a position that is very close to the opposite DNA 5^′^ end, making contact also via Gln-40 and Pro-42 (a schematic of the contacts is provided in Figure S2-a). Also, two hydrogen bonds within a cut-off distance of 0.3 nm are observed between Arg-18 of Zn1 and thymines at position 3 and 4; while Arg-122 of Zn2 makes hydrogen bonds with C41 and C42. Some of these ‘trademark’ interactions were already described in the NMR study of Ref.^14^. Notably, it has been known for quite a long time^36,37^ that the interaction between phosphate groups of DNA and lysine, arginine and phenylalanine residues of TATA-BOX protein helps in producing and stabilizing a sharp kink in the double-helix, and open up the DNA conformation, by inserting two deformed *α*-helices in the minor groove. Similarly here, the PARP-1 molecule makes contact via Gln-150, Arg-122, Arg-18 and Phe-44, found in the Zn1 and Zn2 domains. Moreover, Zn1 digs in the SSB, and reaches for an additional contact with T23 on the opposite DNA strand using its base stacking loop(mainly Met-43 & Phe-44) (Fig. S2-a). All these interactions, along with the trademark interactions, appear to assist and maintain DNA bending and kinking^14,18,38^.

Overall, the average conformation of PARP-1 after the long MD simulation gets back to a structure rather close to the reference experimental one, as far as the contact regions with DNA are concerned. Also, the DNA maintains its strongly bent conformation, as shown from the distribution of bending angle in Figure 3c (black line). The global orientation of Zn1-Zn2 with respect to the DNA is slightly rotated by 17° about the DNA main axis, compared to the experimental reference. However, this has minimal impact on the contacts between the protein and DNA.

#### 3.2.2 Starting from the straight hairpin

The next step was to repeat the docking and MD simulation, but starting from the unperturbed straight conformation of the hairpin, an attempt to mimic the stage of PARP-1 randomly approaching the DNA region containing an SSB. The docking and selection of binding candidates were performed in the same way as above. However, in this case the docking of the whole protein fragment, that is, the two Zn1-Zn2 zinc fingers connected together by a short stretch of amino acids, did give complex results. In the best docked configurations, Zn2 is found in close contact with the 3^′^ DNA end (G45), but Zn1 remains far from interacting. Moreover, the relative position of Zn1 and Zn2 is such that Zn1 lies on the same side as the 3^′^ end, and therefore to get into a proper interaction with the 5^′^ end, as experimentally observed, it would require a large rotation by about 180° of the whole protein about the main DNA axis. Such results could be due to an HADDOCK limitation or an artifact, or could be indicative of a real physical-chemical issue in the interaction with the straight hairpin, which at this stage is difficult to assess. It is worth noting that the recent FRET experiments^18^ using the same DNA hairpin configuration, identified a high affinity for the Zn2 binding alone - albeit with somewhat lower values than for the Zn1-Zn2 complete pair - and vanishing affinity for Zn1 binding alone.

Hence, we selected the best docking configuration having the right orientation of Zn1-Zn2 with respect to the SSB ends, even if this is not the one giving the best HADDOCK score and RMSD value. This configuration was then used as the starting input structure for a long MD simulation of 0.6 *µ*s at T=310K. This starting configuration has Zn2 interacting with G1 (the 5^′^ SSB end, instead of 3^′^) and T23, that is the isolated nucleotide facing the SSB. After the long MD run, however, it evolves into a more correct configuration (Figures 3a and S3-b), with Zn1 approaching the 5^′^ end of the SSB, albeit the closest residues are Ser-41 and Asp-45, rather than experimentally-observed Phe-44 (Ser41 even makes a H-bond with T23). Moreover, Zn2 is penetrating further in the SSB opening, and Leu-151 interacts with T23. On the other hand, Zn2 has a rather weak interaction with the 3^′^ end, via its Pro-149 and Gly-152, and the residue Ile-154 coming within H-bonding distance with G45 for most of the simulation time.

According to our protocol, we performed clustering of the whole trajectory to extract the representative conformations of the interacting system. A comparison of the opening *d*_*g*_ and bending *θ*_*b*_ of the SSB is given in the plots of Figure 3b,c, between the two simulations: (i) starting from the experimental reference (black lines), vs. (ii) starting from the straight hairpin (red lines). After 500ns MD simulation, the DNA appears nearly straight and the gap closed, with little or no room for Zn2 and Zn1 to make contacts with the SSB ends, as it can be observed in the plots in Figure 3b,c. It seems therefore that blind docking is unable to give a proper initial interaction, when the hairpin is not in a already (partially) open configuration. To understand if the initial orientation of Zn1-2 could play a role in the SSB opening and kinking, we therefore proceeded with manually adjusting the orientation of the protein into different interacting configurations, as explained in the next section.

#### 3.2.3 Manually fixing the right DNA-PARP-1 contacts

Given the difficulty of obtaining a starting docked structure corresponding to the experimental observations, we tried to manually approach the Zn1-Zn2 fragment to the DNA hairpin, by using CHIMERA manipulation tools. We initially positioned Leu-151 from Zn2 close to the G45 nucleotide at the 3^′^ end, while the Phe-44 was placed somewhat close to the G1 nucleotide at the 5^′^ end of the SSB.

Starting from this configuration, we performed 150 ps of thermal equilibration, followed by a MD run of 200 ns. The best cluster, shown in the Figure 4a, has RMSD = 0.542 (cutoff = 0.65). This initial configuration resulted in a somewhat bent DNA, closer to what observed in the experiments, along with an increased average distance between the 3^′^ and 5^′^ ends. In Figure 3b,c above, the blue histograms show the bending angle is evolving between *θ*_*b*_=130-170°, while the gap remains open between *d*_*g*_=2-2.5 nm. In addition, Figure 4b displays the plot of distances between reference atoms methyl-C of Leu-151 vs. C2 of G45 (black line); and methyl-C of Met-43 vs. C4^′^ of G1 (red line). The molecular dynamics (MD) simulation reveals that the movement of Zn2 adjusts over time, fluctuating between 0.5-1 nm by the 3^′^. However, Zn1 moves away from the 5^′^ strand after about 100 ns, and fluctuates at a distance between 2-3 nm (see red line in the Figure 4b). Such a significant evolutionary transition of Zn2 and Zn1 could be due to the presence of Arg-122 and Arg-18, which clamp to the major/minor groove of DNA(see again Fig. 4a), bringing about a slight bending in the DNA, and starting to approach the “final” configuration seen in the experimental results.

**Figure 4.**
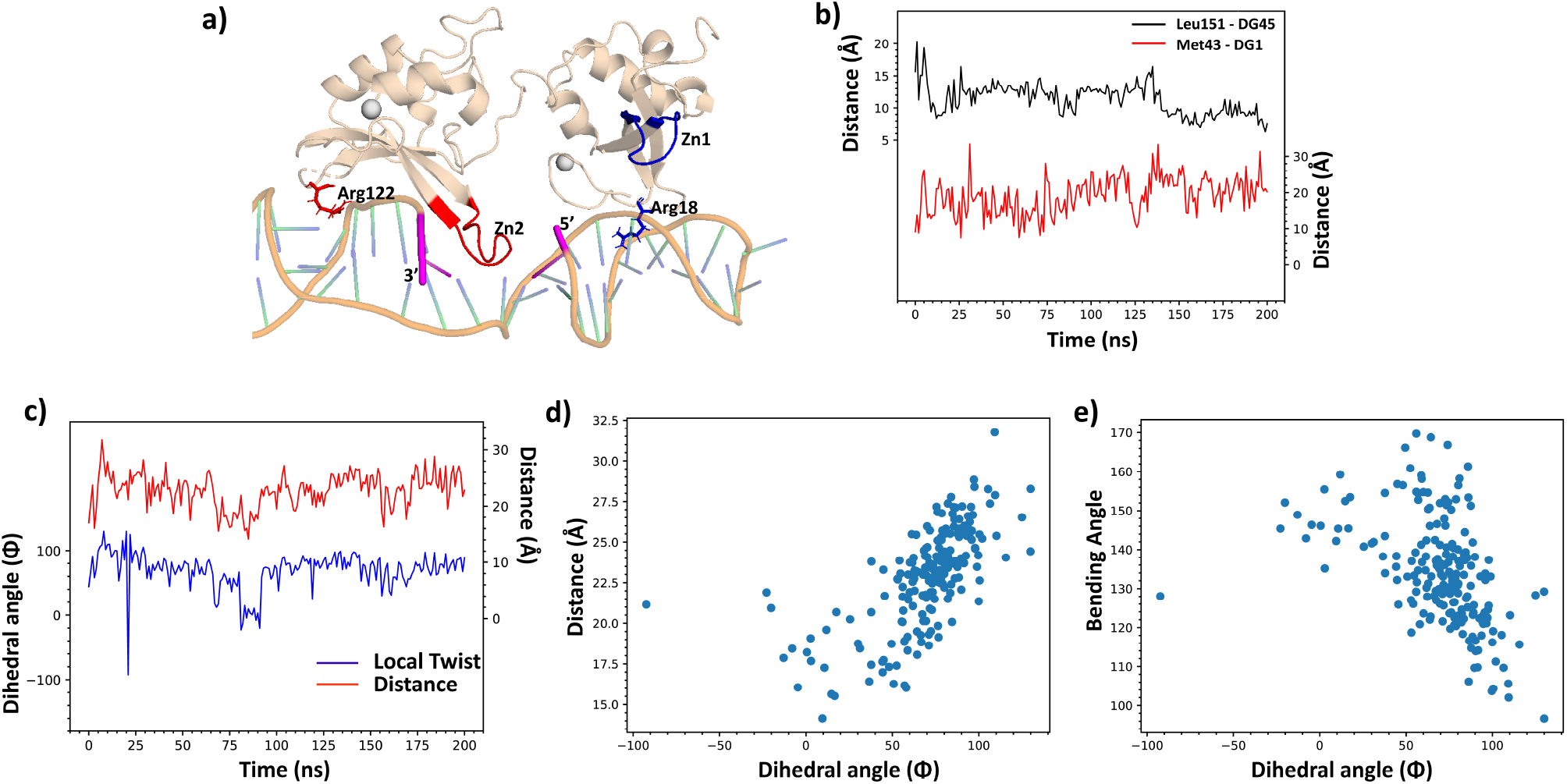
a) Best cluster of the manually-adjusted structure after MD simulation of 200 ns. b) Plot of the C-C reference distances as a function of MD time, for the Leu151-G45 contact at 3^′^ (black line) and the Met43-G1 contact at 5^′^ (red line). c) Plot comparing the local twist and the distance between the 3^′^-5^′^ terminals. d) Scatter plot comparing the evolution of distance between the 3^′^-5^′^ terminals w.r.t local twist. e)Scatter plot comparing the evolution of bending angle w/r to local twist

The degree of SSB opening may be due to the larger or smaller bending angle of the DNA hairpin. However, the distance *d*_*g*_ could increase also because of the local undertwisting of the major groove with respect to the DNA main axis, or because of a combination of bend and twist. To understand this point, we measured the local twist, defined as the dihedral angle Φ formed by the *P* atoms of the four nucleotides around the SSB (1,22,24,45), with large values of Φ corresponding to local DNA undertwisting (Fig. 4c). We also observed a roughly linear correlation between Φ and *d*_*g*_, the increase in the former (undertwisting) resulting in a larger gap (Figure 4d). On the other hand, the correlation plot of Φ vs. *θ*_*b*_ (Figure 4e) indicates as well a linear relationship, where smaller bending angles are coupled with larger undertwisting. During the whole simulation the bending angle spans the interval between *θ*_*b*_=130-150°, but certain instances witness a very sharp decrease to *θ*_*b*_=100-120° (see Figure S4). Such an occasional, sharp bending is clearly due to the clamping of the two PARP-1 loops centered at Arg-18 and Arg-122, which fit in the grooves on either side of the SSB, along with the mechanical constraints of DNA due to its peculiar hairpin configuration. Overall, these results show that large openings can be the result of either bending or (under)twisting, or a combination of both effects.

### 3.3 Comparison between monomeric and dimeric PARP-1

To ascertain whether the dimerization of PARP-1, and the consequent *independent* action of Zn1 and Zn2 (in fact a long-standing question in the literature^9,16,39^) could possibly play a role in the sensing of the SSB, we performed a series of docking+MD simulations with the two Zn fingers disjointed. At first, we simply dropped the 13-aa stretch that links the two moieties in the monomer. In this case, we observed that the simple removal of the 13-aa linker from the same configuration used in Section 3.2.2 (direct docking to the straight DNA hairpin) does not affect significantly the PARP-1-SSB contact. After a 200-ns MD simulation, the comparison of RMSD and RMSF between the two simulations (Fig. S3, see with-vs. without-linker) displays only minor differences such as the sharp spikes around atoms 375 and 1099 of the DNA, corresponding to nucleotides T11 and T35 in the loops at the opposite ends of the hairpin.

Then, we tested the independent docking of the two Zn2 and Zn1 fragments to the straight DNA hairpin, either simultaneously or sequentially. The rationale for using again the straight hairpin, was to ascertain whether the Zn fingers have the ability to spot a pristine SSB, at least in this interaction mode.

For the **simultaneous** interaction, we used the ‘three-molecule’ docking option of HADDOCK. Zn1 (active residues Met43, Phe44) and Zn2 (active residues Gln150, Leu151) were freely docked to active sites G1 and G45 of the hairpin, without further specification of the mutual interactions. In practice, however, we could never obtain any meaningful starting structures, since all the best HADDOCK scores corresponded to configurations in which Zn2 gets in the SSB, filling the gap between the 5^′^ and 3^′^ ends, and Zn1 is placed somewhere close but outside the SSB, in a weakly interacting position. This is likely another proof supporting the notion that the still-closed SSB has not enough space to accommodate both Zn fingers, and Zn2 is the highest-affinity species to start the interaction, while Zn1 remains peripheral.

Then, for the **sequential** interaction, we started with docking Zn2 alone against the initially-straight DNA hairpin. Since the relative positioning of Zn1 and Zn2 are now independent, there are many more possibilities for the interaction. For the sake of argument, we focused the attention on two, nearly symmetrical orientations of Zn2 with respect to the SSB, namely: **orientation (1)** resembling the experimentally observed arrangement of monomeric-Zn2 with respect to the SSB, and **orientation (2)** with Zn2 mirror-rotated by about 180° about the vertical direction (perpendicular to the main DNA axis). For each case (see the following subsections), firstly an equilibration of Zn2 will be studied, and secondly Zn1 will be added, based on the experimental affinity data from which it consistently appears that Zn2 is the PARP-1 moiety that establishes the strongest contact.

#### 3.3.1 Dimeric PARP-1: the “experimental” Zn2 orientation

For the **orientation (1)**, the MD equilibration for 400 ns at T=310K shows that with this initial conformation, Zn2 has a considerable reactivity towards both the 3^′^ and the 5^′^ ends. As shown in Figure 5a, Zn2 is initially close to 3^′^; however, after about 300 ns of simulation it begins approaching also 5^′^, and for the remainder of the simulation it maintains a steady distance of about ∼4 nm from either DNA ends. At the same time, the C4^′^-C4^′^ distance *d*_*g*_ (SSB gap, blue plot in the Figure 5a), starting from values as large as 2-2.5 nm, settles to a much shorter *d*_*g*_ ∼1 nm at about the same time, indicating the rapid closing of the gap, and with a conformation of the hairpin that turns more and more straightened, compared to the largely bent initial state.

We then performed several distinct HADDOCK docking experiments, in which Zn1 was approached to the 5^′^ end of this pristine Zn2-DNA complex at different time frames of the MD simulation. In the following, we show two representative situations taken at times *t*=2 ns (Figure 5b) and *t*=400 ns (Figure 5c), respectively characterized by a very open, vs. a rather closed hairpin instantaneous conformation. After either initial docking, two 500-ns MD equilibration runs were performed. Detailed bonding and solvation free-energy information was deduced from the analysis of the best cluster for each simulation.

**Figure 5.**
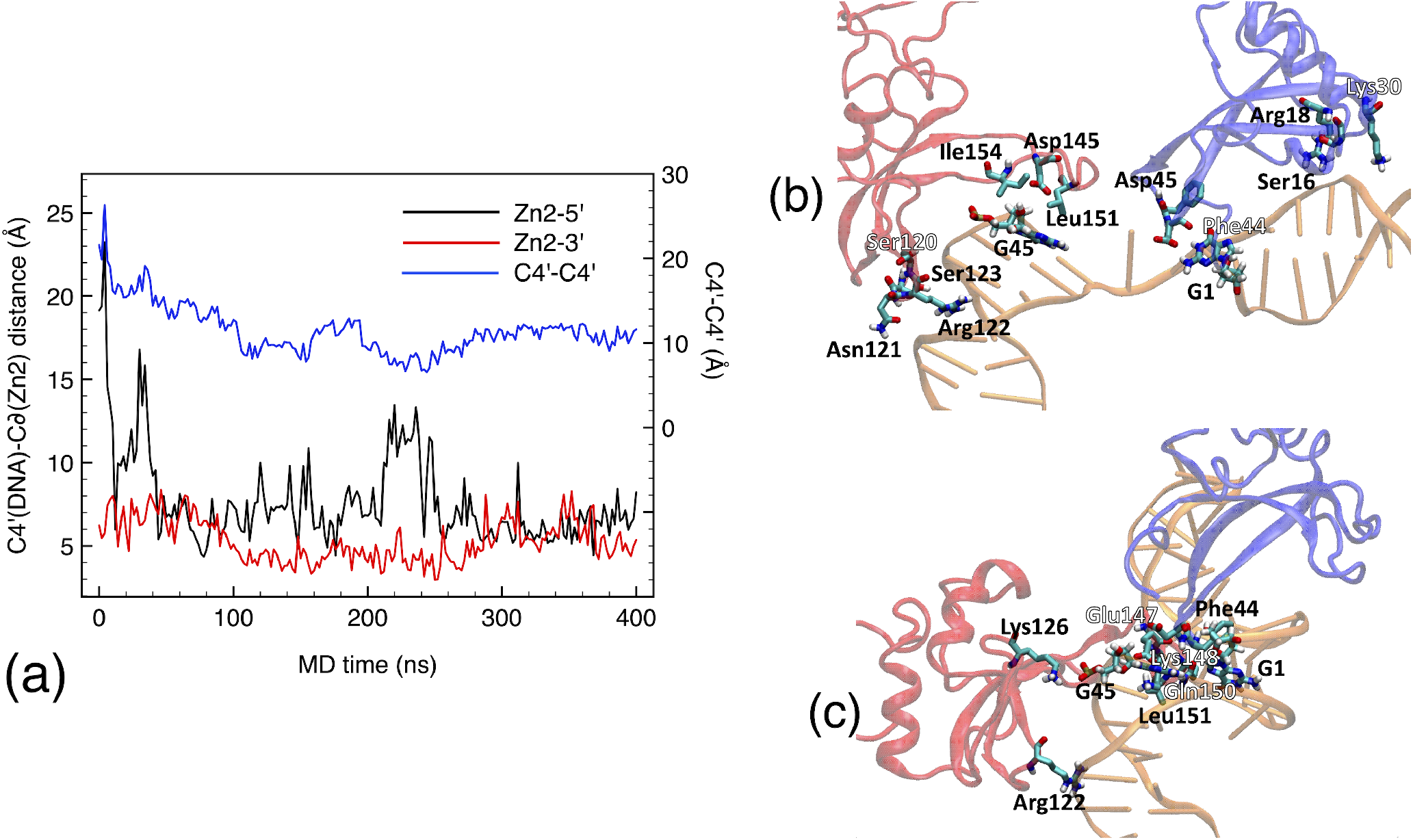
(a) Interaction of Zn2 with the pre-bent DNA hairpin starting from orientation (1, see text). The black and red plots (left Y-axis) represent the time evolution of the distance between the C*δ* of Leu151, and respectively, the two C4^′^ carbon atoms of the 3^′^ DNA terminal (black), or 5^′^ terminal (red); the blue plot (right Y-axis) represents the C4^′^-C4^′^ distance indicating the opening width of the SSB gap. (b) Initial configuration of Zn1 (blue), Zn2 (red) and DNA (orange), after independent docking and subsequent relaxation to the open DNA hairpin (t=2ns). (c) Same as (b), for the (ii) closed hairpin (t=400ns).

For the **t=2ns** docking, the SSB gap always remains quite open at about *d*_*g*_=2 nm, as shown in Figure 6a (black plot), and the bending angle remains constantly about *θ*_*b*_=130° (red plot). The MD trajectory shows a moderate interaction of Zn2 with the 3^′^ DNA end. Leu-151 and Gly-152 still contribute about half of the total hydrophobic contact with G1. However, the total Δ*G*=-2.2 kcal/mol is in large part contributed by the residual interaction of Lys-148 with the middle-exposed DNA base T23, between which a H-bond at 0.195 nm is also formed (Figure 6b, blue plot). Zn1 now appears to interact more strongly with the 5^′^ end, compared to the starting configuration. Phe-44 still maintains a strong hydrophobic interaction, where the phenyl group is mostly facing the G45 heterocycles, and Lys-47 makes an 0.198 nm side-H-bond with the ribose O5^′^. The total (solvation) + (H-bond) Δ*G* is -6 kcal/mol. The direct Zn1-Zn2 interaction is maintained by a strong pair of salt bridges, formed by the N*ζ* nitrogen of Lys-148 and the two carboxyl oxygens of Asp-45. This interaction is likely to distract Zn2 from a possibly closer interaction with the DNA, by holding it closer to the Zn1 loop, rather than to the 3^′^ end.

**Figure 6.**
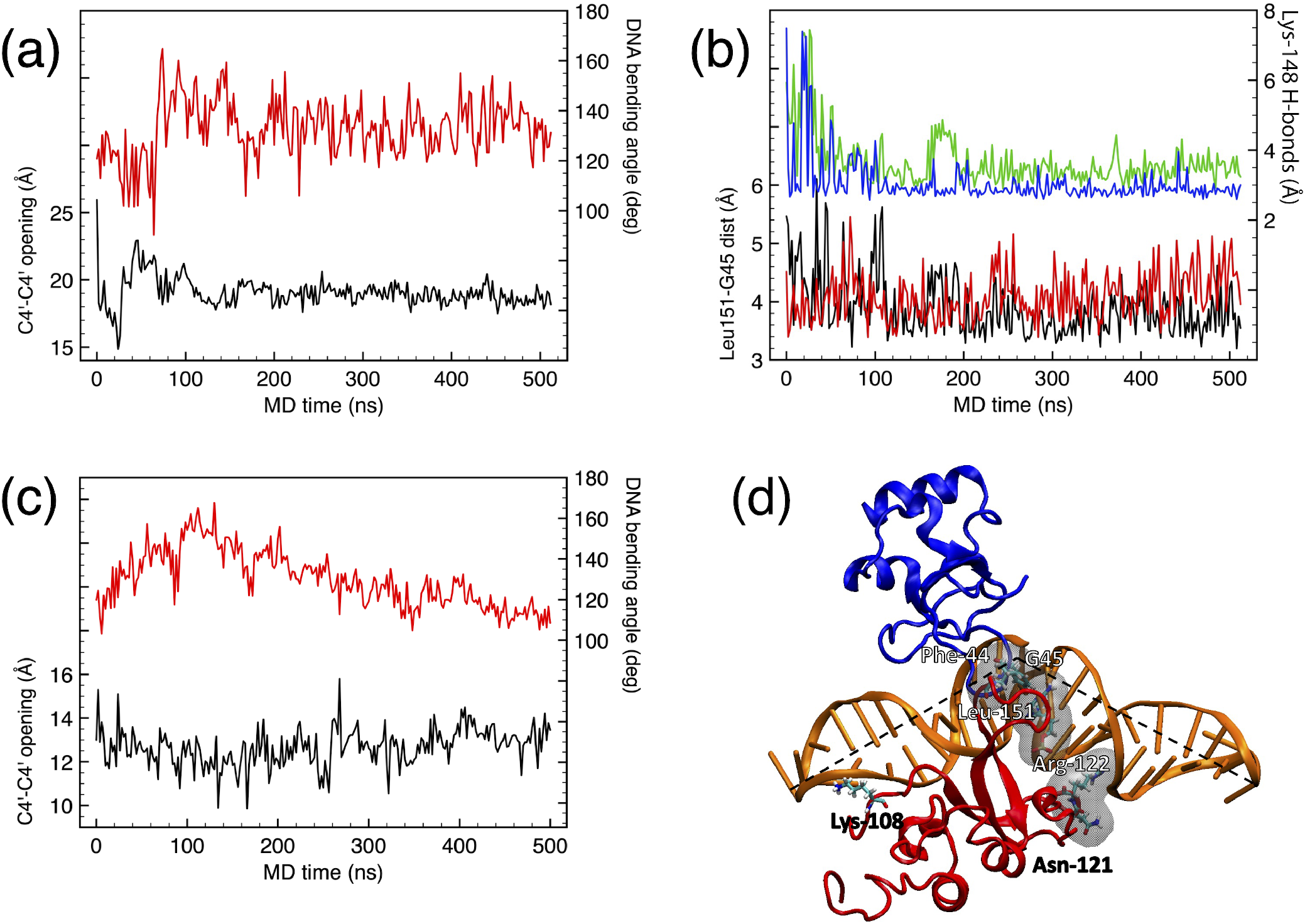
(a) Evolution of the SSB gap opening (black plot, left Y-axis) and of the DNA bending angle (red plot, right Y-axis) for the so-called experimental orientation (see text). (b) Interaction of Zn2 with the 3^′^ DNA end: distance between the two C*δ* of Leu-151 and the C6 (black) and C8 (red) of guanine-45; H-bonds between Lys-148 Nζ and thymine-23 O8 (blue) and N3 (green). (c) Evolution of the SSB gap opening (black plot, left Y-axis) and of the DNA bending angle (red plot, right Y-axis) for the (1-ii) conformation. (d) Best cluster for the (ii) conformation, represented as ribbons (DNA=orange, Zn1=blue, Zn2=red); the interaction regions are depicted as shaded surfaces; the dashed line represents the instantaneous DNA bending angle.

Concerning the **t=400ns** docking, the initial arrangement of Zn1 and Zn2 appears to remain little altered for the whole 500ns MD; because of their mutual interaction, both zinc fingers seem to have a reduced interaction with the SSB gap. However, an interesting information obtained from the plots of this simulation in Figure 6c includes the DNA bending angle rapidly evolving towards quite smaller angles, about 120° in the second part of the trajectory, while at the same time the SSB gap remains quite closed at 12-13 nm. Given the little interaction of both Zn1 and Zn2 with the SSB gap, and the contact of Asn-121 and Arg-122 apparent in Figure 6d (already in slight contact also at the beginning of the simulation, see Figure 5c), such a bending appears to be the result of spontaneous thermal fluctuation. In fact, only at much later times a third contact (Lys-108, see figure) will arise on the opposite side of the DNA hairpin, stabilizing the opening angle.

#### 3.3.2 Dimeric PARP-1: the 180°-flipped Zn2 orientation

For the **orientation (2)** of Zn2 with respect to the straight hairpin, the best docking pose resulted in a cluster with an RMSD 0.128 nm, all the poses of the cluster having the base loop of a.a. 149-152 of Zn2 very close to the 3^′^ DNA end. After 400-ns MD equilibration of the top-scoring pose, a conformation with a large opening of the SSB gap was obtained, with Zn2 reaching its stable position along the 3^′^ end. While this conformation approaches an opening angle similar to that observed in the experiments, it should be equally possible that the kink is due to a spontaneous fluctuation of the DNA hairpin, and/or by direct action of the Zn2 finger penetrating in the SSB gap. By comparing the *θ*_*b*_ bending angle of the phosphate backbone, and the SSB opening *d*_*g*_, it is observed that the DNA gradually unfolds with *θ*_*b*_ fluctuating between 120-130° and *d*_*g*_=20-25 nm.

The average structure obtained as the best cluster from the MD simulations was then used as the starting Zn2-DNA complex, to further dock-in the Zn1 monomer. An initial configuration with Zn1 docked to the SSB gap was obtained from HADDOCK. This combined Zn1-Zn2-DNA configuration is therefore our best candidate for further equilibration at finite temperature and pressure. After 1 *µ*s of MD simulation, the trajectory is clustered and representative configurations were extracted as usual.

It is found that Zn2 penetrates in the SSB gap and has a total Δ*G*=-3.9 kcal/mol, which mainly includes: (a) interaction with the G45 at 3^′^ end;(b) hydrophobic interaction of the 119-122 a.a. loop; (c) Arg-122 making two H-bonds with the O2 oxygen of C19 (0.211 nm); (d) O4^′^ ribose oxygen of C20 (0.249 nm) at the minor groove; and (e) contact with C24 and G25 at the opposite side of the SSB.

Concerning Zn1, Figure 7 shows a summary of the results of MD simulation, focusing on the special binding configuration of Zn1 at the 5^′^ end (Zn2 removed from the view for the sake of clarity). Zn1 strongly interacts with G1, by making a strong H-bond of Pro-49 with the ribose O5^′^ and a weaker one with C4^′^ (Figure 7b), plus a very peculiar contact of Phe-44 (see below). At a later time it also establishes a strong contact with the DNA groove, the Arg-18 making two H-bonds with the O6 oxygen of G4 and G6. Its overall Δ*G* at this later stage is close to -8 kcal/mol, largely coming from the 43-45 a.a. loop and the 5^′^ terminal.

**Figure 7.**
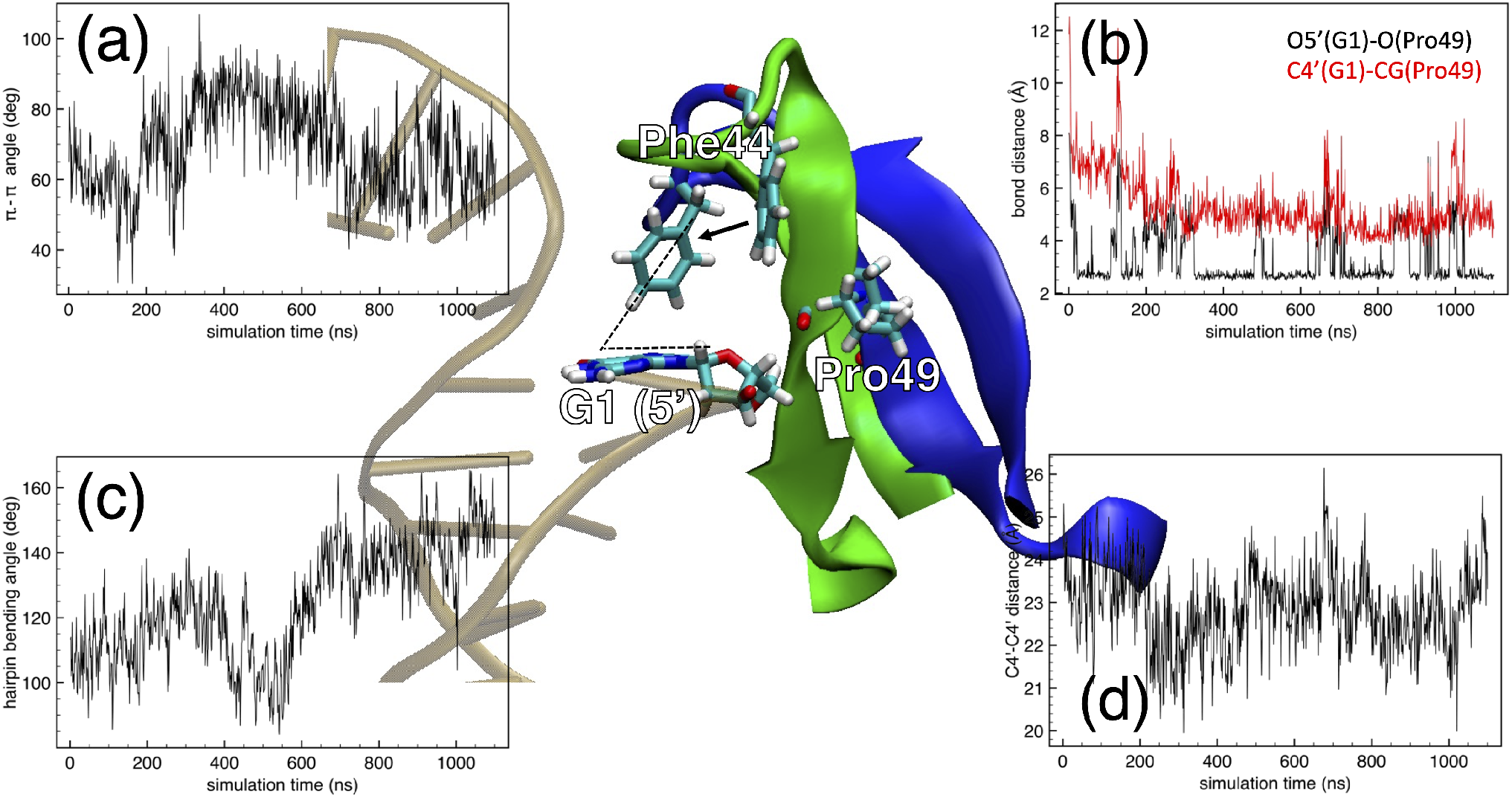
Interaction of the Zn1 with the DNA 5’ terminal after 1.2 *µ*s (blue ribbon), starting from the manually docked configuration (green ribbon). Only the contact loop is depicted. Part of the DNA sketched as a brown ladder in the background. (a) Time plot of the *π* −*π* angle between the planes containing the aromatic cycles of Phe44 and G1 (sketched by the dashed angle in the central panel); the hexagonal ring of Phe44 moves from the initial to the final position as indicated by the little black arrow. (b) H-bonding distances between G1 and Pro49. (c) DNA hairpin bending angle, measured between the P atoms of nucleotides 12-24-34. (d) Opening of the 5^′^-3^′^ gap in the DNA hairpin, measured by the distance between the respective C4^′^ atoms.

The interaction of Phe-44 with the aromatic cycles of guanine G1 is of a special “edge-to-face”, or T-type, which was shown to be one characteristic feature of protein-DNA interactions^40^. In such *π*-bonding configuration, the phenyl of Phe-44 makes an angle with the plane of the G heterocycles typically between 60-70°, rather than coupling in a side-by-side stacked arrangement. Phenylalanine is found to be the most likely candidate for such T-type bonding with DNA, twice more frequently with thymine compared to purines^40^. Such a bonding geometry for our case is shown in the central panel of Figure 7, where the *π*-*π* angle is also defined. The evolution of the Phe44-G1 angle is shown in Figure 7a, which fluctuates between 60 and 80°, a figure in good agreement with the literature findings^40^.

By looking at Figure 7c,d there seem to be no apparent correlation between the fluctuations of the T-bonding of Zn1, and either the DNA bending angle (Figure 7c), or the SSB gap opening (Figure 7d). The former evolves from a rather bent shape in the first ∼500ns, with *θ*_*b*_=90-120°, to a more closed shape in the second half of the MD trajectory, with *θ*_*b*_=140-160°. However, the relative arrangement of Zn1-Zn2 is such that the SSB gap (Figure 7d) remains always very open, at *d*_*g*_=2.2-2.4 nm, independently on the straightening of the hairpin.

Clearly, such results for orientation (2) are rather interesting when compared to those obtained with the orientation (1), albeit in this (2) the Zn2-Zn1 orientation is someway “opposite” to the (1) (instead closer to experimental observations). They seem to suggest that, if a dimer made of two copies of PARP-1 can independently interact with the SSB (a possibility not yet fully explored in the experiments), a wider range of interaction possibilities arises. Of course, some of these configurations would be excluded in the realistic situation with two full copies of PARP-1 interacting, instead of using just the Zn fingers, since the whole proteins could lead to steric clashes of the dimer. Overall, the ensemble of simulations of this Section 3.3 points out that the role of Zn2 in pre-determining the opening of DNA is relevant, but might not be as crucial as early suggested. It is rather the particular structure and sequence of the DNA hairpin, which possibly allows the SSB gap to open up spontaneously by frequent, free thermal fluctuations. Only when the fluctuation spontaneously opens up the hairpin, Zn2 could make its interaction, stabilize the DNA kink, and subsequently make room for Zn1. As shown, this can also occur with mutual arrangements quite different from those observed in experiments in which monomeric PARP-1 is crystallized.

## 4 Discussion

To summarize, in the present work we investigated by all-atom molecular dynamics simulations the competition-cooperation of two Zn-finger domains of PARP-1, in recognizing a single-strand break (SSB) in a model fragment of DNA. Since our main purpose was to make a link with recent structural biology experiments^14,18^, we used their same 45-nucleotide ssDNA hairpin designed with a sequence such that the structure makes two near-symmetric loops and closes back with a missing base between the 5^′^ and 3^′^ terminations (the guanine-1 and guanine-45, facing the central, unpaired nucleotide thymine-23). Both in the published experiments and in our simulations, the gap between the two terminal bases was taken as representative of a fully-formed SSB with proper 5^′^-dRP and 3^′^-OH terminations, to which we also added a second configuration with both OH-terminated ends, for the sake of comparison. Such situations can be representative of an enzymatically-cut SSB or the final stage of processing radiation-induced SSBs, just prior to the start of the repair phase. The PARP-1 protein was represented in full atomic resolution only for the regions actively used in the co-crystallization experiments, specifically, the two Zn fingers Zn1 and Zn2. In some of the MD simulations we kept them physically connected by their 13-a.a. linker, while in other simulations we made them act independently to simulate a PARP-1 dimer.

We used a combination of molecular docking with empirical free-energy functionals, and molecular dynamics simulations with all-atom interatomic force fields, to investigate different stages of the Zn1-Zn2 identification of the SSB. We explored configurations ranging from the structure with the DNA hairpin straight and the gap “closed”, to the experimentally identified configurations in which the hairpin is strongly bent and the SSB gap is wide open. The main findings of our study can be summarized as follows:

1. The highly peculiar hairpin conformation of the DNA structure allows for a large conformational variability under thermal fluctuations at physiological temperature, compared to straight, long DNA harboring a similar SSB at its center. While the straight DNA can experience broad curvatures, especially when its length approaches and surpasses the persistence length (≳50 nm), the atomic structure surrounding the SSB remains relatively unaffected. It is therefore possible that, also in the experiments, the DNA hairpin could spontaneously open up to extreme deformations (such as, bending angles much smaller than 120-130°, SSB gap openings of 2 nm or more), thanks to its special conformation and even without a direct help from the PARP-1 contacts.
2. The experimental structure of the interacting complex^14,18^ can be readily retrieved in the MD simulations, starting from the separate fragments initially taken in the respective experimental conformations. This means that the experimental interacting structure represents indeed a global thermodynamic minimum of the DNA-Zn1-Zn2 complex also for the molecular simulation, thereby providing a strong support to the reliability of the simulation protocol, and of the subsequent results.
3. The interaction of PARP-1 with the DNA hairpin in an initially straight conformation is not able to display the direct transition to the “final” state, as represented by the experimental structure. We observed that Zn2 is indeed able to force itself inside the SSB gap and make contact with the 3^′^ end, attaining values of bending angle and gap opening not far from the experimental. However, the position of Zn1 in these instances is not correct and lies far from the 5^′^ DNA end, unlike what is seen in the experiments. A series of MD simulations in which both Zn1 and Zn2 were forcefully placed in a better initial position with respect to the straight SSB, demonstrated a steady contact between Zn1-5^′^ and Zn2-3^′^. However, in this case the SSB gap does not open and the hairpin remains practically straight (i.e., the DNA-protein contact remains “external” to the SSB). It cannot be excluded that such a configuration could possibly lead to a final state closer to the experimental one, if observed over a time scale much longer than that accessible to MD simulations.
4. Resistance to large DNA bending is apparently due to the extra contacts that both Zn fingers may establish, by pushing their side loops (the a.a. 17-19 for Zn1, and the a.a 120-122 for Zn2) deep in the DNA grooves flanking the SSB gap, which forbid further bending of the hairpin. It may anyway be speculated that, over a much longer time scale than the *µ*s, such ‘side contacts’ could eventually lead to a forced bending. Overall, we observed a good correlation of the SSB gap opening with the DNA bending angle, as well as with the degree of local undertwisting of the double helix. From a kinetics point of view, it appears that the DNA bending is mostly due to spontaneous, free thermal fluctuations until, at some later moment, the side protein loops contact a DNA groove, and “lock-in” the conformation, be it a straight or a bent hairpin. On the other hand, the SSB gap opening is initially correlated with the DNA bending and twisting, but it is ultimately determined by the arrangement of the Zn fingers, primarily Zn2 which has the tendency to penetrate deep in the gap, and interacts with the nucleotides opposite to the SSB.
5. The question whether PARP-1 operates as a monomer, using the Zn1 and Zn2 from the same molecule for detecting the SSB, or as a dimer, cooperatively using Zn1 and Zn2 from two different molecules, remains an open one. We performed computational experiments by using opposite arrangements of the two Zn fingers, either acting in parallel or sequentially. We find that Zn2 displays the highest affinity for the SSB open ends; it attacks preferentially the 3^′^ end, coherently with the experimental findings, but it can also attack both ends at the same time, especially if starting from a relatively more closed SSB gap. The role of Zn1 is therefore secondary, as shown in the experiments (which however used only monomeric PARP-1). Also in the dimeric form, Zn1 must fight its way to the closed SSB, and it is only when the SSB is already substantially open, that it may join Zn2 in recognizing the SSB. Notably, on occasion we also observed conformations in which Zn1 bridges the two loops of the hairpin, forcing extreme bending angles, for Zn2 to interact with the SSB.

By putting together the findings of Section 3.2.3 about PARP-1 interaction with the straight and closed SSB, next to the observations of Section 3.2.1 about the easier flexibility of the DNA hairpin interacting with the Zn2 alone, and the observations of Section 3.1, about the large opening and bending fluctuations observed for the isolated hairpin in the absence of PARP-1, an interesting interpretation of the dynamics of recognition of the SSB by PARP-1 can be reconstructed. A possible picture that emerges from the present work, in agreement with the *final* structures found in experiments, can be that the Zn2 first interacts with the 3^′^ end of the SSB, thereby helping the spontaneous DNA bending; after which, Zn1 can start interacting with the 5^′^ end, leading to the final observed kinking. The fact that the direct docking of the Zn1-Zn2 couple on the pristine closed SSB gives a good interaction, as far as the adhesion free energy, but with little or no opening of the SSB, because of the extra contacts along the DNA sides, adds support to the need for spontaneous thermal fluctuations of the DNA hairpin. The main objection to such a picture is, why then the experiments can observe Zn1-Zn2 interacting with a wide-open SSB, and a largely bent DNA hairpin? The answer can be found in the free fluctuations of the isolated hairpin *prior* to interaction: it is possible that a spontaneously fluctuating DNA, offering every now and then the SSB in a wide-open conformation, has a larger affinity for Zn1-Zn2 compared to Zn2 alone, and this is what has been interpreted as Zn1-Zn2 forcefully opening the SSB (the so-called ‘bind-then-bend’ action). Notably, from the preliminary comparison of the interaction of PARP-1 with a free DNA strand, it appears that this mode of interaction is not entirely specific to the peculiar construction of the hairpin. While in the hairpin the terminal loops greatly facilitate ample structural oscillations, also the free DNA shows enough bending upon contacting the protein, in this case associated with its long-wavelength fluctuations. An interesting way to prove experimentally this idea could be to try to use a hairpin construct but with much longer flanking sequences, so as to keep the closing loops as far as possible, left and right of the SSB gap, thus approaching the free DNA condition.

As far as future work with MD simulations, it will be our next priority to translate the free-DNA-PARP-1 interaction in the context of linker DNA joining nucleosomes. Interestingly, in that case the free fluctuations of the DNA will be severely constrained by the presence of massive nucleosomes at both ends, whose relative displacements can however induce further mechanical deformations by coupling bending, twisting and kinking modes^41^. Moreover, it should be considered that, at least in radiation-induced SSBs, the gap terminations are far from clean 5^′^*/*3^′^ ends at the stage of initial defect detection. Therefore, the ability of PARP-1 to recognize a “dirty” terminated SSB remains another key issue to be elucidated.

## Acknowledgements

The authors acknowledge funding from the University of Lille Project PEARL “Senesimex: experiments and modelling of radiation-induced cell senescence”. FC also acknowledges partial funding from the ANR Project “Dyprosome: dynamics of DNA repair proteins at nucleosomes”. We thank generous computing time allocation on the JEAN-ZAY supercomputer of IDRIS-CNRS in Orsay, under projects GENCI A0130712986 and A0150712986.

## Author contributions statement

F.C. and P.A.S. conceived the computer experiments. P.A.S. conducted the molecular dynamics simulations. All authors analysed the results and reviewed the manuscript.

## Additional information

Partial trajectories and input files for the molecular dynamics simulations in this work are available at the repository FigShare, accession link: 10.6084/m9.figshare.25957288. Additional information available from the corr. author (.

## Confilcts of interest

The authors declare no competing interests.

## SUPPLEMENTARY MATERIALS

**Figure S1.**
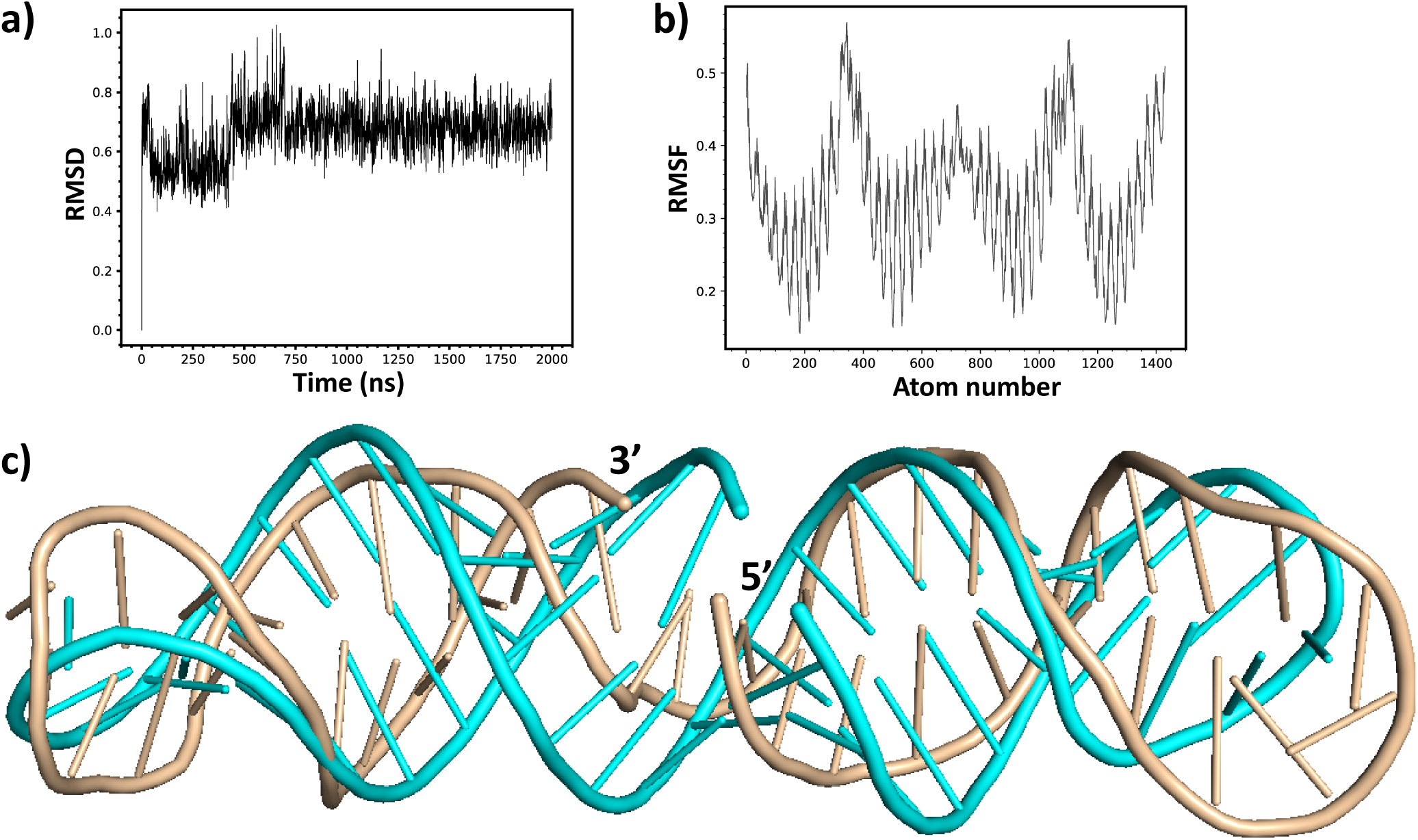
(a) RMSD values of isolated DNA hairpin in configuration **1** over 2*µ*s MD simulation. (b) RMSF value on a residue-by-residue basis. (c) Superimposed structure of the hairpin loop best cluster after 2µs simulation (gold), and the initial structure (cyan).

**Figure S2.**
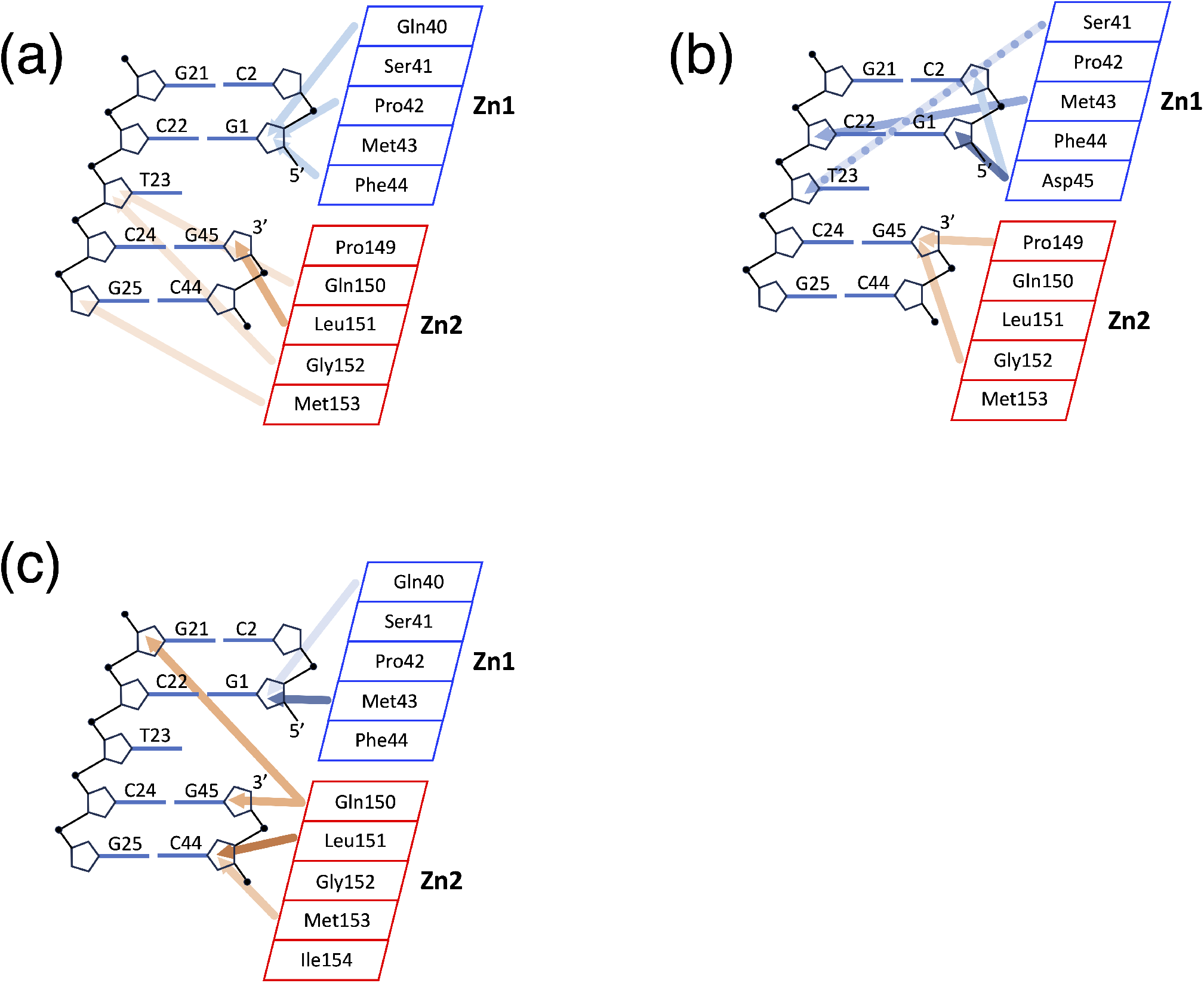
Contact maps of the PARP-1-DNA hairpin assembly (Zn1 blue, Zn2 red) including an SSB (missing nucleotide at position 46). Arrow color intensity according to increasingly close bonds. (a) Docking + MD of PARP-1 Zn1 and Zn2 domains against a DNA hairpin in the NMR experimental configuration **1**. (b) Docking + MD PARP-1 Zn1 and Zn2 domains against the straight DNA hairpin configuration **2**. (c) After MD of PARP-1 manually fitted by CHIMERA to the straight DNA hairpin **2**.

**Figure S3.**
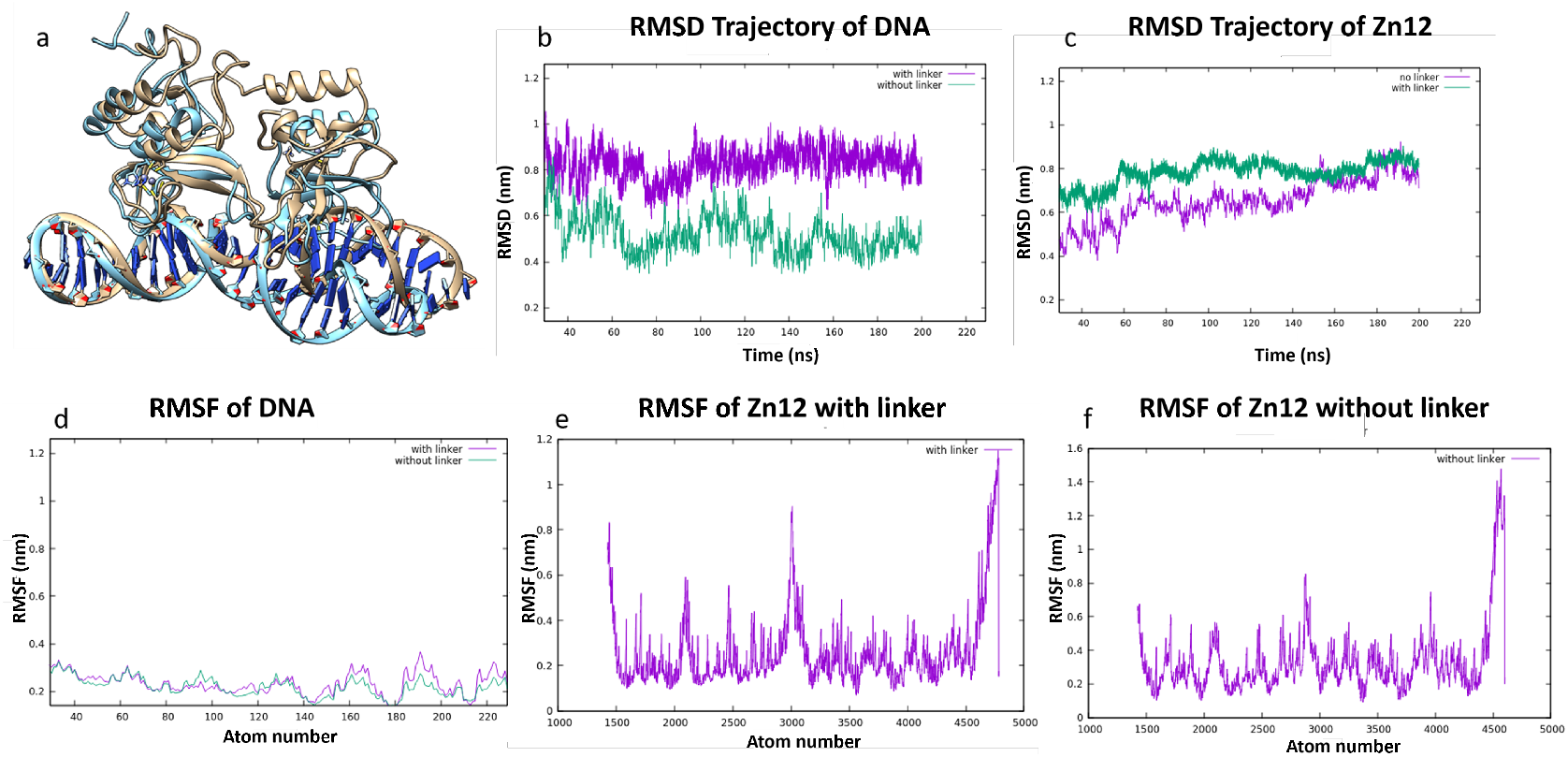
(a) Superimposed structure of the best cluster of Zn1-Zn2 interacting with DNA after 200ns-MD simulation, with (gold) or without (cyan) the 13-aa linker joining the zinc fingers. (b) RMSD trajectories of DNA, and (c) of Zn1-Zn2, purple= with linker, green= without linker. (d) RMSF of DNA by residue, with and without linker. (e,f) RMSF of Zn1-Zn2 with/without linker.

**Figure S4.**
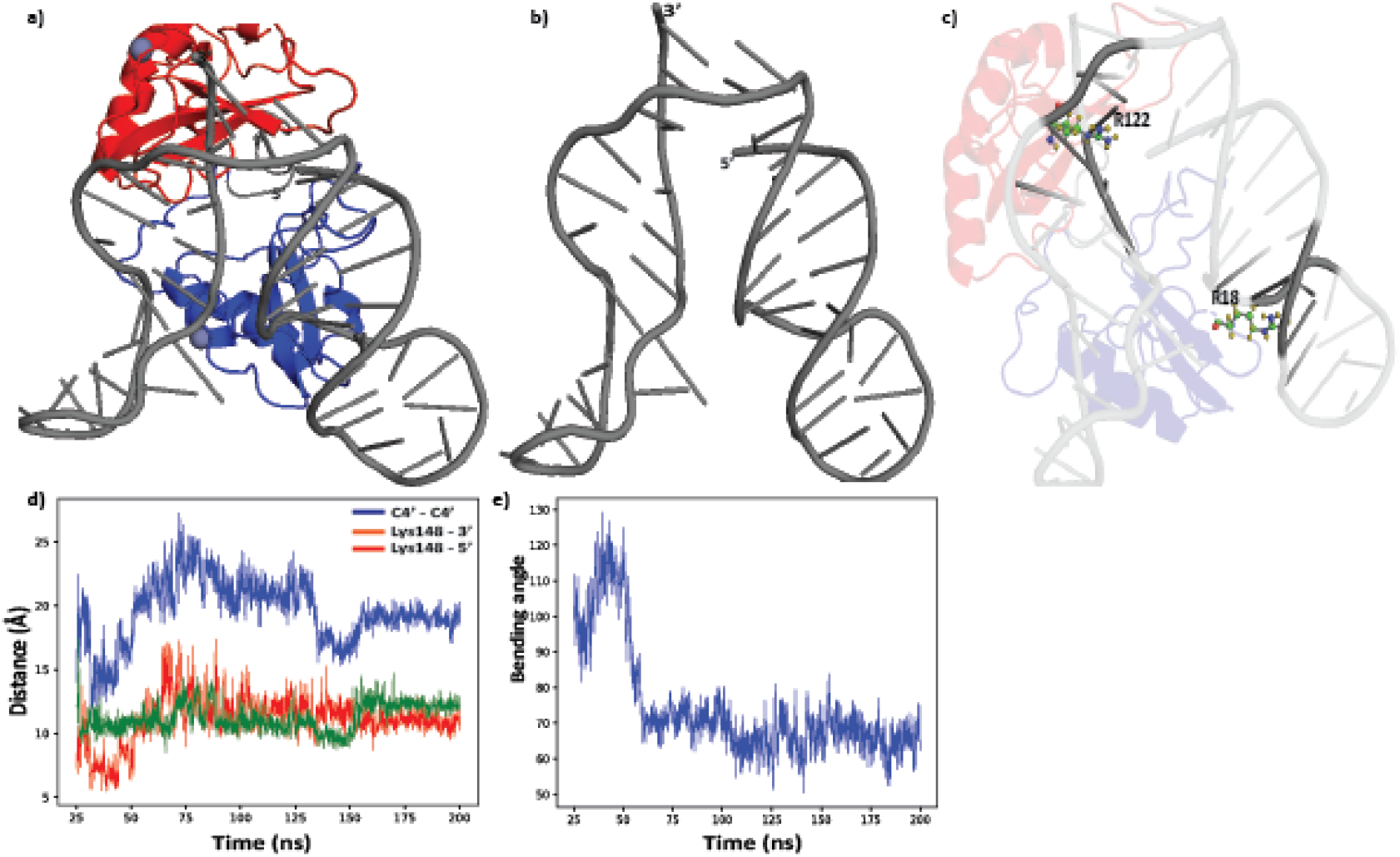
(a) Best cluster obtained after 200ns-MD simulation with manually adjusted Zn1-Zn2 starting from the straight hairpin. Notice Zn2 (red) interacting with both SSB ends, and Zn1 (blue) locking the bent DNA conformation. (b) Snapshot of DNA bending during the simulation. c) Snapshot of Arg-18 and Arg-122 penetrating into the minor groove of DNA and imposing the bending. d) Evolution of SSB gap opening (blue, C4^′^-C4^′^ distance); red,green=distance between Zn2’s Lys-148 and the 5^′^ or 3^′^ ends throughout the 200ns simulation. (e) Evolution of DNA bending angle over the same 200ns.

## Notes

### Competing Interest Statement

The authors have declared no competing interest.

